# Does Vergence Affect Perceived Size?

**DOI:** 10.1101/2020.02.23.961649

**Authors:** Paul Linton

**Affiliations:** Centre for Applied Vision Research, City, University of London, Northampton Square, Clerkenwell, London EC1V 0HB

**Keywords:** Visual Scale, Size Constancy, Vergence, Taylor Illusion, Multisensory Integration

## Abstract

Since Kepler (1604) and Descartes (1637), it’s been suggested that ‘vergence’ (the angular rotation of the eyes) plays a key role in size constancy. However, this has never been tested divorced from confounding cues such as changes in the retinal image. In our experiment participants viewed a target which grew or shrank over 5 seconds. At the same time the fixation distance specified by vergence was reduced from 50cm to 25cm. The question was whether the reduction in the viewing distance specified by vergence biased the participants’ judgements of whether the target grew or shrank? We found no evidence of any bias, and therefore no evidence that eye movements affect perceived size. If this is correct, then this finding has three implications: First, perceived size is much more reliant on cognitive influences than previously thought. This is consistent with the argument that visual scale is purely cognitive in nature (Linton, 2017; 2018). Second, it leads us to question whether the vergence modulation of V1 contributes to size constancy. Third, given the interaction between vergence, proprioception, and the retinal image in the Taylor illusion, it leads us to ask whether this cognitive approach could also be applied to multisensory integration.

## Introduction

As objects move forwards or backwards in space, the image they cast on the retina varies drastically in size. And yet objects don’t appear to change dramatically in size when they move closer or further away. This suggests that there is a neural mechanism (‘size constancy’) that compensates for the drastic changes in the retinal image caused by changes in distance (for a review see Sperandio & Chouinard, 2015). We can distinguish between two kinds of visual cues for size constancy: 1. Pictorial Cues, which are present in the static monocular retinal image (such as familiar size and perspective), and which account for the impression of size constancy in pictures. And 2. Triangulation Cues, which typically rely on introducing multiple viewpoints either simultaneously (vergence and binocular disparities) or consecutively (motion parallax).

1. Pictorial Cues: The neural correlates of size constancy are better understood for pictorial cues since 2D pictures are more easily presented to participants in fMRI scanners (e.g. Murray et al., 2006). However, pictorial cues are neither necessary nor sufficient for size constancy. First, pictorial cues are unnecessary because, as we shall discuss below, observers in the Taylor illusion appear to experience something close to full size constancy from vergence alone. Second, pictorial cues are insufficient because, as Sperandio et al. (2012) observe, size constancy in pictures is merely a fraction (10%-45%) of the size constancy experienced in the real world (Murray et al., 2006; Leibowitz et al., 1969; Goodale, 2020 estimates 10%-30%; see also Millard et al., 2020 for a recent attempt to disambiguate static monocular and binocular size constancy, although recognising vergence may have influenced their monocular results). To spell out this claim, consider the kind of stimulus used in Murray et al. (2006):

We judge the further ball in Fig. 1 to be many times larger in physical size than the nearer ball. But we also judge the further ball to take up more of the picture than the nearer ball even though they have the same angular size in the picture. This apparent increase in angular size is the phenomenon that size constancy is concerned with. In Murray et al. (2006) the further ball was judged to 17% larger in angular size than the nearer ball.

**Figure 1.**
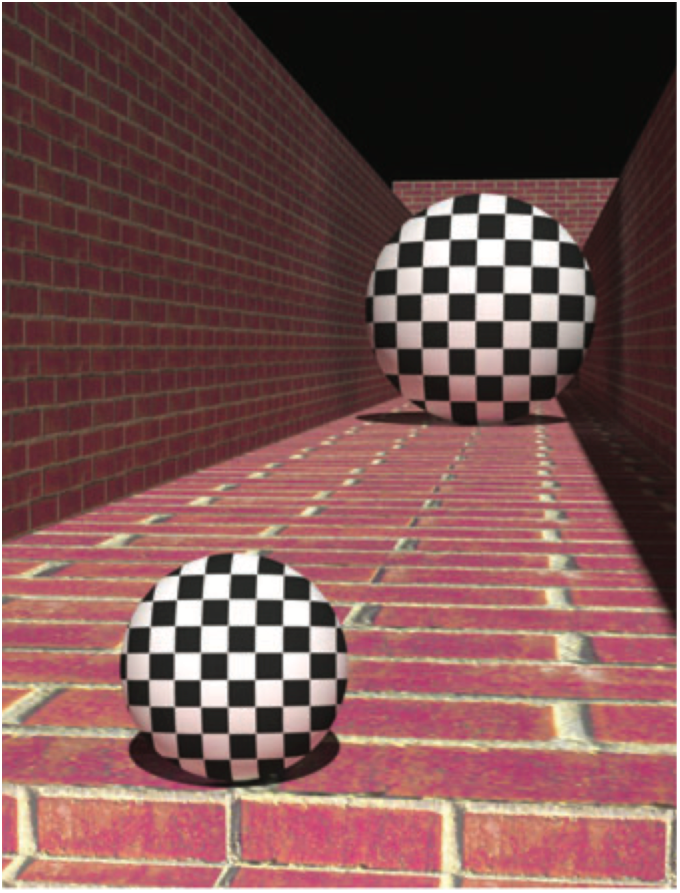
Demonstration of pictorial size constancy in the stimulus used by Murray et al. (2006) (© Scott Murray)

When reviewing the literature, we have to keep the distinction between perceived physical size and perceived angular size in mind. For instance, Sperandio et al. (2012) contrast the partial (10%-45%) size constancy in pictures with the full (100%) size constancy experienced in the real world according to Emmert’s Law (Emmert, 1881). However, Emmert’s Law is a claim about perceived physical size not perceived angular size. Indeed, the essence of Emmert’s Law is the claim that perceived physical size varies proportionately with perceived physical distance when the perceived angular size is fixed. By contrast, an angular size interpretation of Emmert’s Law would falsely imply that objects do not reduce in perceived angular size with distance.

Nonetheless, even with the distinction between physical size and angular size in mind, vergence is thought to play an important contribution to the perceived angular size of objects, known as vergence size constancy.

2. Triangulation Cues: In terms of triangulation cues to size constancy (vergence, accommodation, binocular disparity, motion parallax, defocus blur), the emphasis has been on vergence. First, motion parallax is neither necessary for size constancy (full size constancy is observed by participants in an fMRI scanner in Sperandio et al., 2012), nor is there particularly strong evidence that motion parallax contributes to size constancy (Combe & Wexler, 2010 only marginally qualify “the common notion that size constancy emerges as a result of retinal and vergence processing alone”). Second, binocular disparity is typically regarded as merely providing relative depth, and not absolute size and distance. Still, considering size constancy only requires relative changes in distance to be matched by relative changes in apparent size (distance_1_ ÷ distance_2_ = size_1_ ÷ size_2_), a merely relative depth cue could suffice. The problem is that binocular disparity doesn’t even provide relative depth information until it has been scaled by absolute distance information, which is typically assumed to come from vergence. As Brenner & van Damme (1998) observe, a 1° change in retinal disparity could equally reflect a change in distance of 20cm to 21cm (5% increase in distance) or 2m to 4m (100% increase in distance).

There is also a deeper conceptual point. Although Linton (2017; 2018) explores the possibility that vision may be divorced from absolute size and distance, orthodox discussions of size constancy typically articulate it in terms of the visual system using absolute distance to determine perceived size. For instance, Emmert’s Law (Emmert, 1881) is typically articulated as S = c(R x D), where S is perceived size, c is a constant, R is retinal image size, and D is perceived distance. But, as we already mentioned, binocular disparity is typically thought of as being a merely relative depth cue outside very limited circumstances (objects taking up at least 20° of the visual field; Rogers & Bradshaw, 1995). Instead, vergence is typically cited as being one of our most important absolute distance cues at near distances (Mon-Williams & Tresilian, 1999; Viguier et al., 2001; for a review and challenge to this consensus, see Linton, 2020).

Kepler (1604) and Descartes (1637) were the first to suggest that the visual system uses vergence (the angular rotation of the eyes) to scale the size of the retinal image. Evidence for vergence size constancy has come from four specific contexts where it has been found that changing the vergence angle affects the perceived size of objects (so-called ‘vergence micropsia’):

1. Wallpaper Illusion: Before the invention of the stereoscope by Wheatstone (1838), the earliest evidence of vergence micropsia was the ‘wallpaper illusion’, the observation that if you cross your eyes whilst looking at a recurring wallpaper pattern, the wallpaper pattern appears smaller and closer (Smith, 1738; Priestley, 1772; Goethe, 1810; Meyer, 1842, 1852; Brewster, 1844; Locke, 1849; Lie, 1965; Ono et al., 1971; Kohly & Ono, 2002; see Howard, 2012 for review).
2. Stereoscopic Viewing: The invention of the stereoscope by Wheatstone (1838) (which presents separate images to each eye) enabled the eyes to be rotated independently of the retinal image. Wheatstone observed that if eye rotation was increased, the perceived image appeared to shrink, even though the images shown to each eye remained fixed (Wheatstone, 1852; Helmholtz, 1866, p.313; Judd, 1897; Frank, 1930; Hermans, 1937, 1954; Locke, 1938; Adams, 1955; Von Holst, 1955a, 1955b, 1957; Heinemann et al., 1959; Gogel, 1962; Biersdorf et al., 1963; Wallach & Zuckerman, 1963; McCready, 1965; Leibowitz & Moore, 1966; Leibowitz et al., 1972; Komoda & Ono, 1974; Regan et al., 1986; Enright, 1989).
3. Telestereoscopic Viewing: Building on Wheatstone (1838), Helmholtz (1857) invented the telestereoscope, and observed that if we use mirrors to artificially increase the distance between the two eyes, the world appears miniaturised. In his Treatise on Physiological Optics he observed that “it will seem as if the observer were not looking at the natural landscape itself, but a very exquisite and exact model of it, reduced in scale” (Helmholtz, 1866, p.312). This effect has been attributed to vergence by Helmholtz (1857; 1858; 1866, p.310) and Rogers (2009; 2011), since the eyes need to rotate more to fixate on the same physical distance (cf. Linton, 2018 for an alternative account discussed below), and has been extensively studied in the military research (where helicopter pilots often view the world through cameras with increased interpupillary separation, see Newman & Ostler, 2009; Stuart et al., 2009; and Priot et al., 2010; 2011; 2012; 2018).
4. Taylor Illusion: Vergence is also thought to be central to the multisensory integration of hand motion and the retinal image in the Taylor illusion. If you make an after-image of your hand with a bright flash, and then in complete darkness move your hand closer to your face, the after-image of your hand appears to shrink even though it is fixed on the retina (Taylor, 1941). The best current explanation for the Taylor illusion is that it is due (Taylor, 1941; Morrison & Whiteside, 1984; Mon-Williams et al., 1997) or almost entirely due (Sperandio et al., 2013) to the increase in vergence as the eyes track the physical hand in darkness (see also Gregory et al., 1959; Carey & Allan, 1996; Bross, 2000; Ramsay et al., 2007; Faivre et al., 2017a; and for vergence scaling of after-images see Urist, 1959; Suzuki, 1986; Lou, 2007; Zenkin & Petrov, 2015). Importantly, when Sperandio et al. (2013) moved the participant’s hand and vergence in opposite directions, they found that (a) the after-image size changed in the direction of vergence, not the hand movement, and (b) the magnitude of the size change when vergence and the hand were in conflict was almost as large as when both the hand and vergence were moving in the same direction.

Surveying the literature on vergence micropsia, two things are striking: First, to our knowledge, there has never been a report of a failure of vergence micropsia within peripersonal space (near distances corresponding to arms reach). Even on the rare occasions when a change in vergence fails to provide an impression of motion-in-depth (for instance, when motion-in-depth is vetoed by a stimulus that takes up the whole visual field) as in Regan et al. (1986), the authors still report “apparent size changes as about threefold when convergence changed from about 0 deg to 25 deg”, with the authors observing: “Changes in size and depth produced by ocular vergence changes are well known”.

Second, the after-image literature appears to suggest that vergence provides something close to perfect size constancy for distances between 25-50cm. This can be seen for two reasons: First, because size constancy appears close to perfect for 25-50cm when vergence is the only distance cue. Apparent size doubled for the representative subject in Sperandio et al. (2013) (incongruent condition) from 3.3cm at 25cm (suggested by the *y* = –0.61*x* + 3.3 line of best fit) to 6.3cm at 50cm (average of size estimates after a >3° vergence eye movement) (my analysis of their Fig. 5 using WebPlotDigitizer 4.2; Marin et al., 2017). Second, the same conclusion is arrived at by a combination of the fact that (a) the Taylor illusion provides near perfect size constancy in this distance range (Bross, 2000; Ramsay et al., 2007; Sperandio et al., 2013), coupled with the fact that (b) the Taylor illusion can be attributed almost entirely to vergence (Sperandio et al., 2013).

Vergence size constancy is therefore regarded as a fundamental aspect of visual perception. However, we believe that vergence size constancy should be re-evaluated for two reasons:

First, our recent work suggests that vergence is an ineffective absolute distance cue once confounding cues have been controlled for. Participants are unable to use vergence to judge absolute distance (Linton, 2020), and we are reluctant to embrace the possibility (raised by Ono & Comerford, 1977 and Bishop, 1989) that vergence might still be an effective size constancy cue even if it proves to be ineffective for absolute distance judgements.

Second, one surprising fact is that to the best of our knowledge vergence size constancy has never been tested divorced from confounding cues (changes in the retinal image, such as diplopia or retinal slip, or changes in hand position) which inform the observer about changes in distance. The reason for this is easy to appreciate. Vergence can only be driven in one of two ways. Either participants track the retinal slip of a visual object moving in depth (such as an LED: Mon-Williams et al., 1997; Sperandio et al., 2013), in which case participants are informed about the change in distance by binocular disparity (the apparent retinal slip of the stimulus as it moves in depth). Or participants track their own hand moving in depth (as in the Taylor illusion), but this gives them proprioceptive information about the changing distance instead. The purpose of our experiment was therefore to test vergence size constancy in a context where it is known to be effective (vergence changes over 5 seconds from 25cm to 50cm: Sperandio et al., 2013), but in a way that controls for subjective knowledge about the changing distance.

## Materials and Methods

Participants viewed two targets on a display fixed 160cm away through two metal occluders that ensured that the left eye only saw the right target and the right eye only saw the left target. We were therefore able to manipulate the vergence distance specified by the targets (indicated by the arrow in Fig. 2) by changing the separation between the targets on the fixed display.

**Figure 2.**
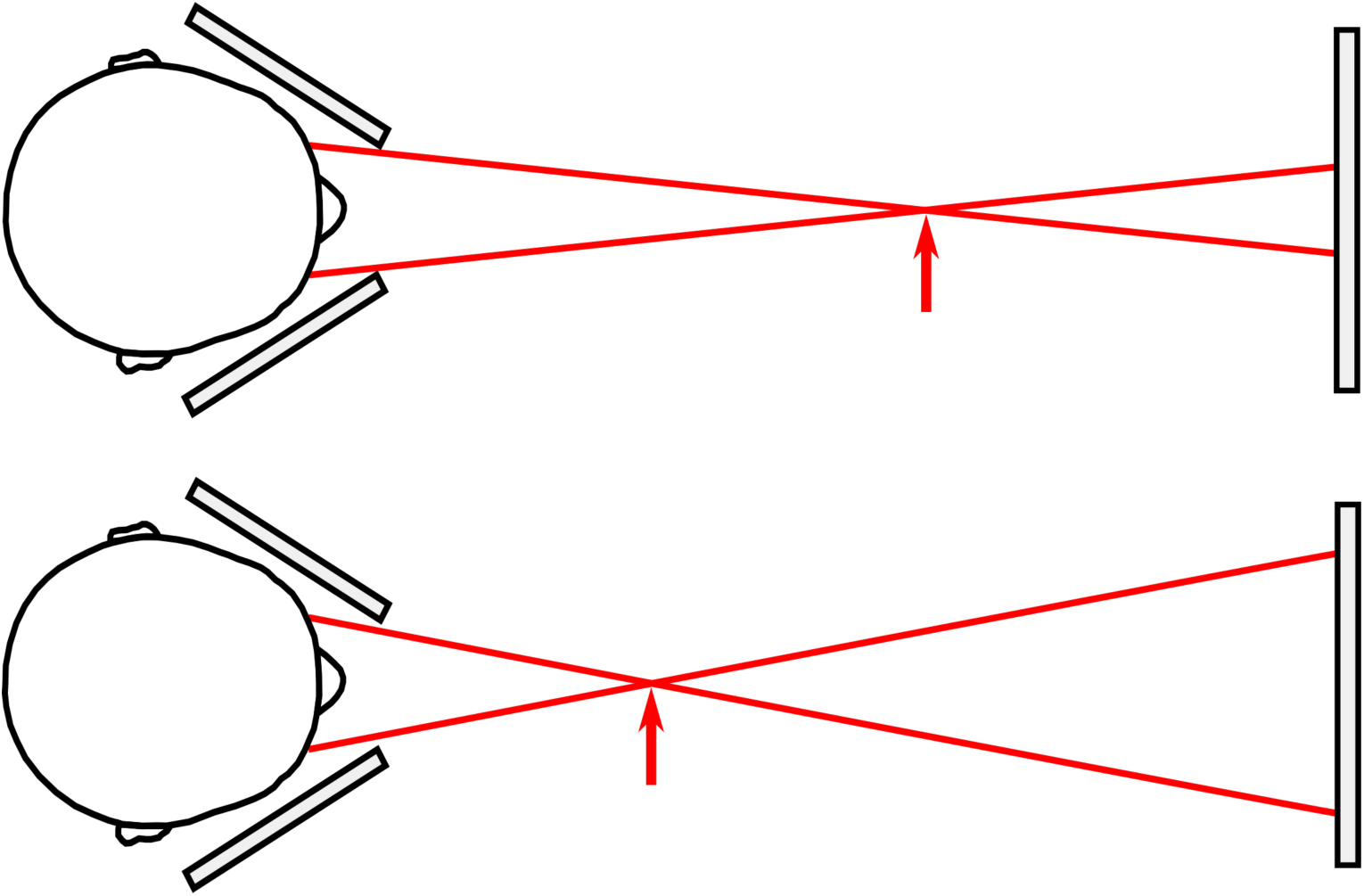
Illustration of how the vergence specified distance was manipulated in the experiment using a fixed display. By increasing the separation between the targets on the display (compare top and bottom images) we were able to reduce the vergence specified distance.

Before the experiment began, participants rotated the metal plates to ensure that they only saw one target in each eye, and were asked to report if the targets ever went double. Our apparatus therefore relied upon (a) manipulating the vergence demand (the vergence specified distance of the stimulus), coupled with (b) a subjective criterion (diplopia; whether the target went double) to ensure the vergence response was effective. No participant reported that they experienced diplopia during the experiment, although (as we discuss below) 1 participant was excluded from the outset because they couldn’t fuse the target, and 2 because they experienced the target as blurry.

Quantifying the effect of vergence on the perceived size of the target posed a complex technical challenge which we resolved in five ways:

First, we built an objective estimate of size change into the target itself. The vergence size constancy literature typically relies upon viewing an after-image (which has a fixed retinal size) and then asking participants to subjectively evaluate the change in perceived size after a vergence change by (1) asking participants to match their visual experience to (a) a visible chart (Bross, 2000; Lou, 2007; Sperandio et al., 2013) or (b) a memorised chart (Ramsay et al., 2007), or (2) by asking participants to make a conscious size judgement of (a) the physical size of the after-image (Mon-Williams et al., 1997), or (b) the after-image’s % size change (Carey & Allan, 1996). We wanted to eradicate this highly cognitive and subjective evaluative process with a more objective criterion of the target size change. If increasing vergence reduces the perceived size of the target then we can quantify this by increasing the physical size of the target during the vergence change and estimating at what increase in physical size participants are at chance in determining whether the target got larger or smaller. The target (illustrated in Fig. 3) consisted of two horizontals bars connected by a vertical bar. It was 3° in angular size (width and height) at the start of the trial. On each trial the vergence specified distance of the target reduced from 50cm to 25cm over 5 seconds. At the same time, the physical size of the target on the display increased or decreased by between −20% and +20%, and participants had to make a forced choice using a button press whether the target got bigger or smaller (“did the target get bigger or smaller?”).

**Figure 3.**
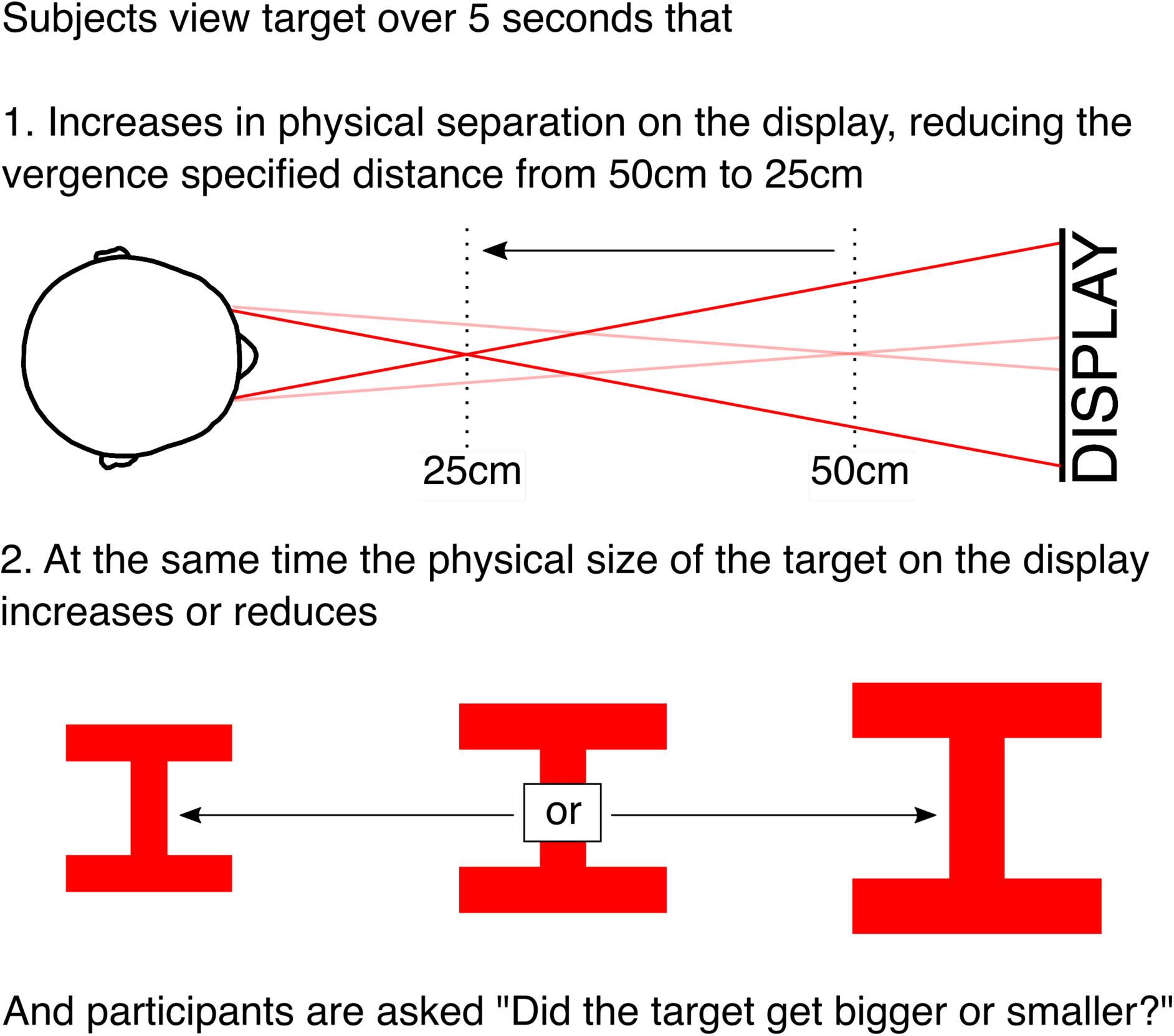
Summary of the two simultaneous manipulations of the targets during each trial of (a) the physical separation of the targets on the display, that manipulated the vergence specified distance of the target, and (b) the physical size of the targets on the display. Participants were asked to judge whether the target got bigger or smaller?

To summarise, over the 5 seconds of each trial we made two simultaneous changes to the targets. First, we changed the physical separation of the targets on the display in order to change the vergence specified distance of the target (Fig. 2). Second, at the same time, we increased or decreased the physical size of the targets on the display by between −20% and +20%.

2. Second, we wanted the target to be presented in darkness to exclude any residual visual cues. An interesting finding from piloting was that the usual technique (having participants view a CRTs through neutral density filters) wasn’t effective in eradicating residual luminance from the display (it degraded the target before residual luminance was completely eradicated). Instead, we achieved this in four ways: First, we used an OLED display (LG OLED55C7V) which, unlike normal displays, does not produce residual luminance for black pixels. Second, subjects wore a mask to block out any residual light, which had red eye filters through which the red stimuli were viewed (blocking out 100% green and ∼90% blue light). Third, subjects viewed the stimuli through a narrow (17°) viewing window of 48cm x 18cm at a distance of 60cm. Fourth, the whole apparatus was covered by blackout fabric, and before the experiment began subjects pulled a hood of blackout fabric over their heads and the external lights were turned off. A photograph and cross-section plans of the apparatus are provided in Fig. 4.
3. Third, in order to drive vergence without providing subjective distance information, we used a visual stimulus that (unlike an LED) provided ‘sub-threshold’ binocular disparities: binocular disparities that are visible to the participant’s visual system (in order to drive vergence), but subjectively invisible to the participant themselves. This we achieved with a 3° target moving in depth from 50cm to 25cm over 5 seconds. Although this motion in depth is gradual (equivalent to an average speed of 5cm/second, corresponding to an average vergence angle change of 1.4°/s), this is consistent with the changes in vergence in Sperandio et al. (2013) (25cm to 50cm over 5 seconds), where close to perfect vergence size constancy was previously reported. We expected that using a target rather than an LED would have this effect for two reasons. First, being slightly larger on the retina, it was likely to improve vergence tracking of the target. Second, any remaining retinal slip would be less discriminable against a slightly larger target.
4. Fourth, in order to present a constant retinal image with eye rotation, we rendered the targets to maintain a constant radius from, and orientation to, the eye. This was achieved in OpenGL by ‘projecting’ the target onto the display, so that the correct retinal image was achieved when the participants viewed the target (Fig. 5) (camera set in OpenGL to nodal point of the eye, and an asymmetric frustum was used so that the far clipping plane matched the distance and dimensions of the display). A bite bar was used to ensure that the nodal point of the eye remained fixed during the experiment (Fig. 4), and the 6mm difference between the nodal point and the centre of rotation of the eye was intentionally ignored (cf. Linton, 2019; Konrad et al., 2019).
5. Fifth, another challenge of this display is that it requires the eyes to focus (or ‘accommodate’) at the distance of the display (160cm), whilst vergence (the angular rotation of the eyes) is at 25-50cm. This decoupling of vergence and accommodation doesn’t happen in normal viewing conditions, and too much vergence-accommodation conflict can lead to the target going blurry or double (Hoffman et al., 2008). To solve this problem we had an optometrist fit each participant with contact lenses (based on the participant’s valid UK prescription) so that the optical distance of the display was 33cm even though its physical distance was 160cm. This ensured a maximum of +/− 1 dioptres of vergence-accommodation conflict, well within the zone of ‘clear single binocular vision’ (Hoffman et al., 2008). Indeed, some of the most dramatic reports of vergence micropsia have been in the presence of large vergence-accommodation conflicts (e.g. 6.5 dioptres in Regan et al., 1986), so the presence of +/− 1 dioptre should not be objectionable.

**Figure 4.**
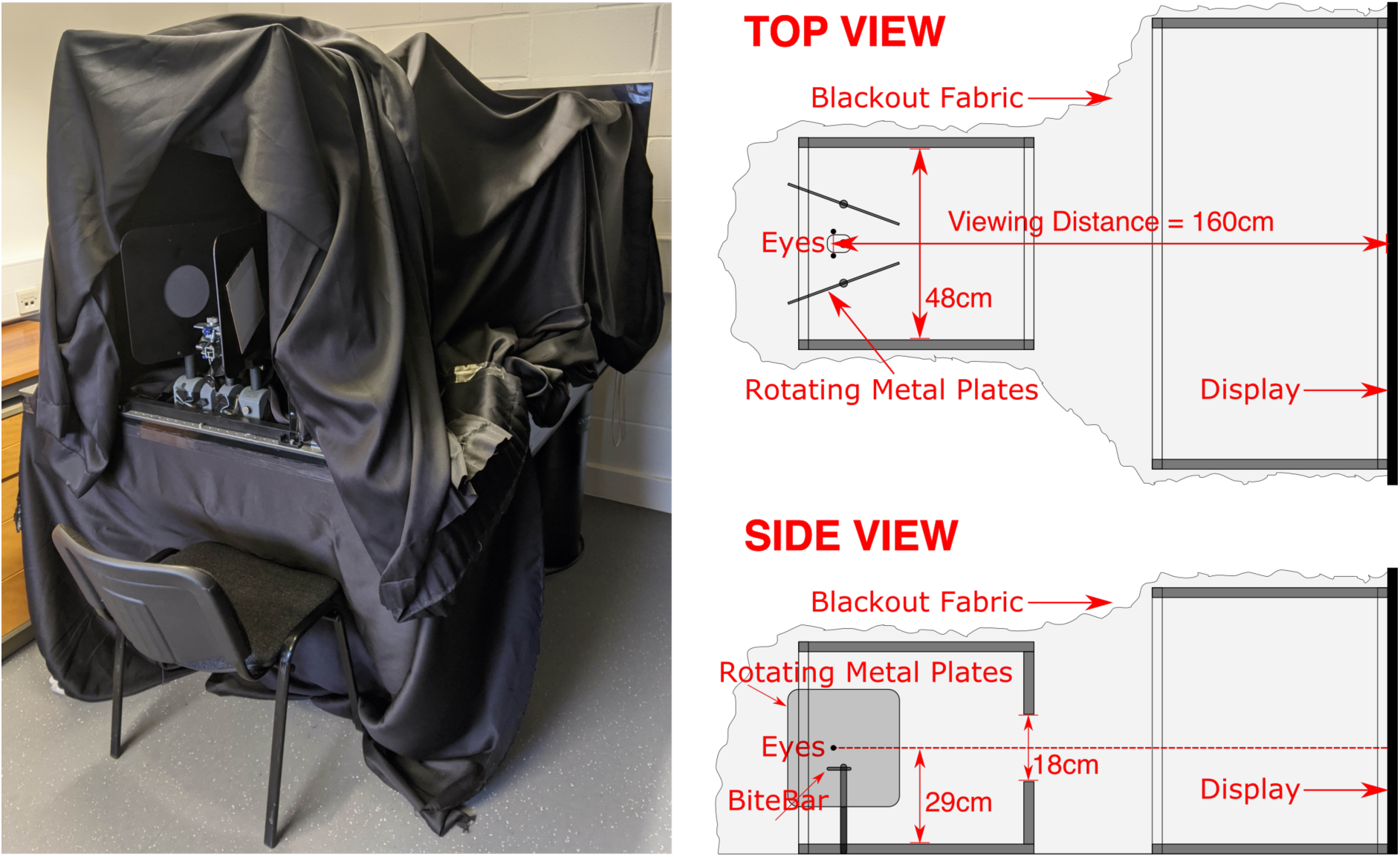
Photograph and cross-section plans of the apparatus.

**Figure 5.**
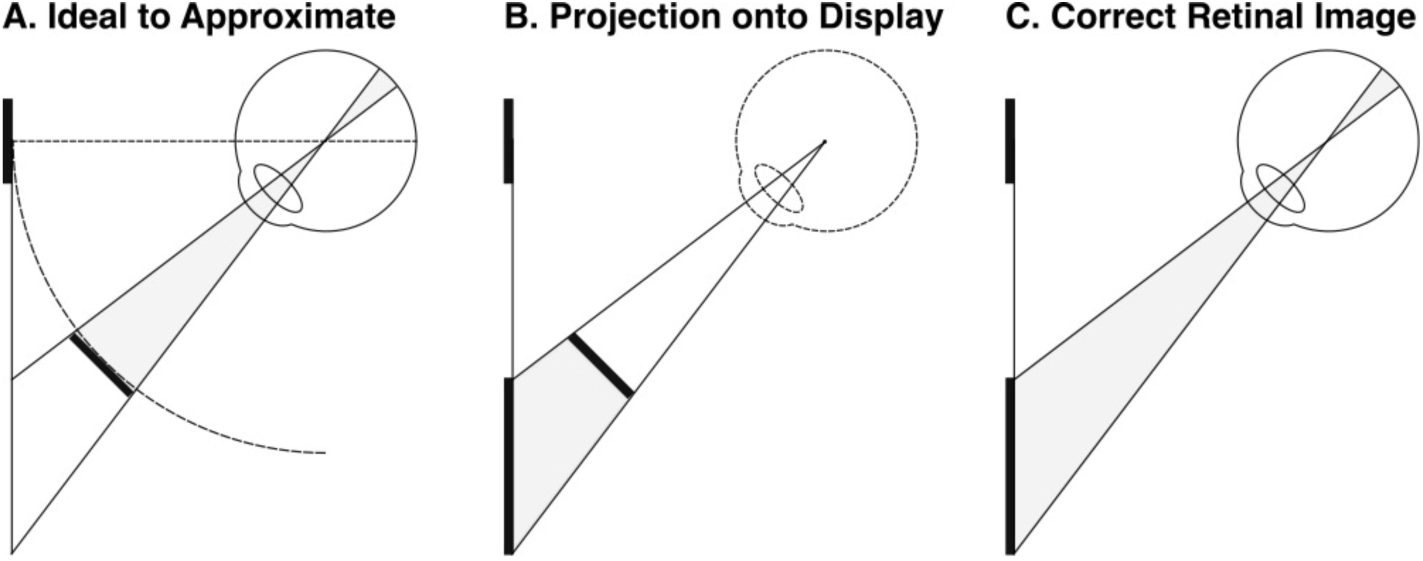
OpenGL rendering of target achieves the correct retinal image for a target with a constant radius and orientation to the eye, whilst presenting the target on a fronto-parallel display.

We used a four-parameter maximum likelihood model (Quest+: Watson, 2017; Brainard, 2017) to estimate when participants were at chance. Participants completed 200 trials (10 sets of 20 trials), and on each trial Quest+ tested the size change that would be most informative. In piloting, we found that the author (an experienced psychophysical observer) could not detect size changes over 5 seconds that were smaller than 1.5%. So, if vergence changes the perceived size of the target by less than 1.5%, vergence size constancy can be dismissed as smaller than the smallest effect size of interest under an inferiority test (Lakens et al., 2018; in our actual experiment this was revised down to 1.43%, the detection threshold of our most sensitive observer).

Assuming vergence doesn’t bias size judgements, how would the number of participants affect our ability to determine this fact? We can simulate the experiment 10,000 times (bias = 0, detection threshold = 5%, lapse rate = 2%), and fit a hierarchical Bayesian model (discussed below) to the data (Fig. 6). We found that with 5 or more observers we should be able to rule out any size constancy effects greater than 1.5%.

**Figure 6.**
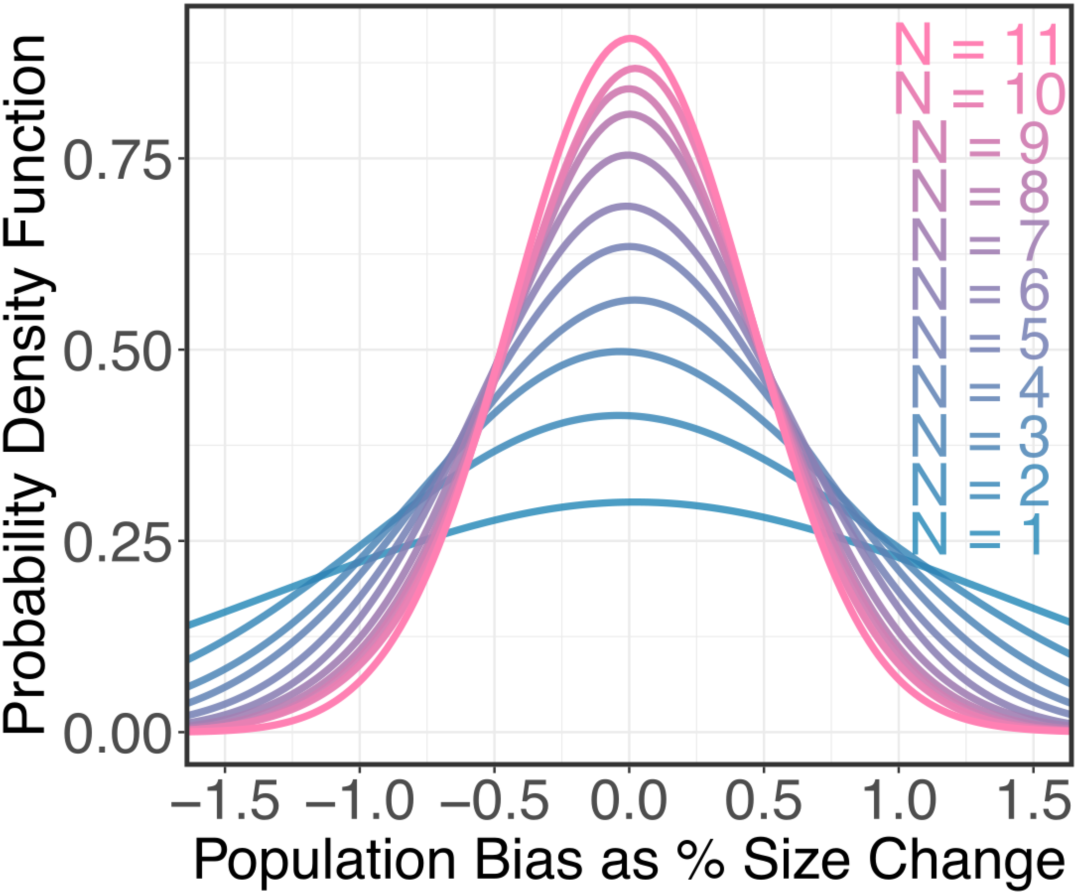
Simulated Experiment. We simulated the experiment 10,000 times in Quest+ (bias = 0, detection threshold = 5%, lapse rate = 2%) to model how increasing the number of participants would improve the accuracy of our hierarchical Bayesian estimate of the true bias (true bias = 0). We determined that we needed n ≥ 5 to rule out an effect greater than our smallest effect size of interest (vergence size constancy > 1.5%).

11 observers (8 female, 3 male; age ranges 20-34, average age 24.5) participated in the experiment: the author and 10 participants recruited using an online advertisement (13 were originally recruited, but 1 was excluded because they could not fuse the target, and 2 were excluded because their vision was blurry with the contact lenses). All participants were screened to ensure their accommodation was within normal bounds for their age (tested with a RAF near-point rule), vergence within normal bounds (18D or above on a Clement Clarke prism bar), and stereoacuity within normal bounds (60 arc secs or less on a TNO stereo test). The author’s participation was required to (a) confirm Quest+ mirrored the pilot data, and (b) provide a criterion for the minimum effect size. All other subjects were naïve as to the purpose of the experiment, and were paid £15/hr for 3 hours. The study was approved by the School of Health Sciences Research Ethics Committee at City, University of London in accordance with the Declaration of Helsinki.

The code for running the experiment is openly available: https://osf.io/5nwaz/, running on Matlab 2019a (MathWorks) with PsychToolBox 3.0.15 (Kleiner et al., 2007).

## Results

Let us consider what we would expect to find according to (a) the null hypothesis (vergence has no effect on perceived size) and (b) the alternative hypothesis (vergence has an effect on perceived size). As we have already discussed, if vergence has no effect on perceived size, then participants should be at chance at determining whether the target got bigger or smaller when we don’t introduce a size change (bias = 0). By contrast, if participants experience something close to full size constancy, then we would have to increase the size of the target by 100% in order to cancel out the reduction in perceived size caused by vergence micropsia (which would equate to a 50% reduction in size, because halving the distance leads to a doubling of the retinal image size, assuming the small angle approximation).

These two hypotheses, and various intermediate degrees of size constancy between these two extremes, are plotted in Fig. 4A. What the hierarchical Bayesian model of our results in Fig. 4A show is that the bias we found was ≈ 0, consistent with there being no vergence size constancy.

To explain our conclusion, the individual results are plotted in Fig. 7C. Each blue dot represents a physical size change that was tested by the Quest+ maximum likelihood model, and the darkness of the dot indicates the number of times it was tested. It’s important to keep in mind that with Quest+ no one data point should be viewed in isolation. The underlying assumption is that Quest+ is fitting a psychometric function to the data. On each trial Quest+ tests the physical size change that would be most informative to the four parameters of the psychometric function being estimated (the slope, the bias, floor, and ceiling). If Quest+ tests one physical size change a few times and finds performance at 100%, and an almost identical size change and finds performance at 0, the interpretation is that the actual response is somewhere between these two extremes. Put simply, from a Bayesian modelling perspective, there’s nothing to suggest that these changes between neighbouring data points reflects a wild variation in participant response.

**Figure 7.**
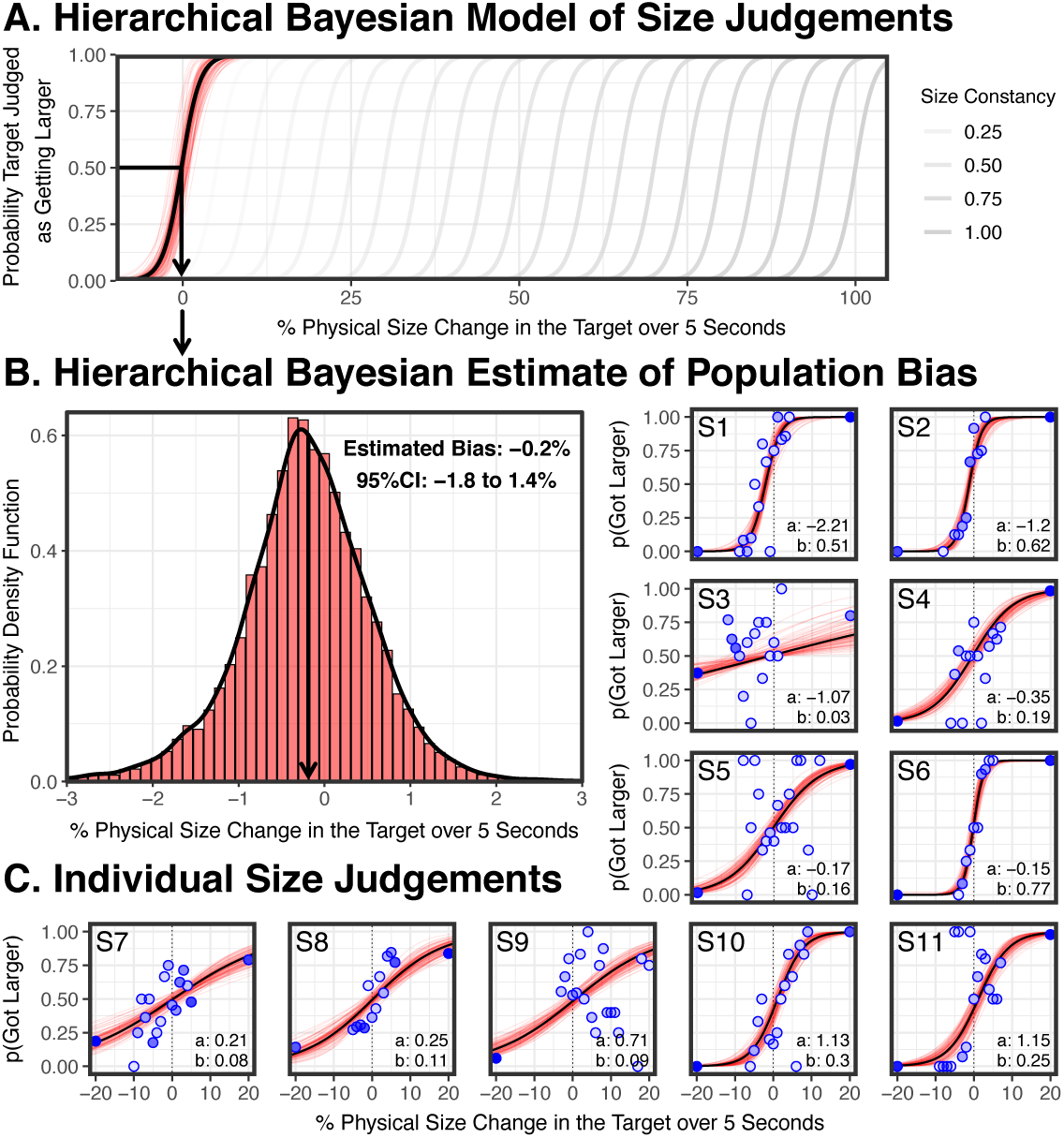
Results. A. Hierarchical Bayesian model of the population psychometric function in black (based on 15,000 posterior estimates, 100 representative posterior estimates in red). Also shown are predictions for various degrees of vergence size constancy effect sizes (in grey). B. Probability density function of 15,000 posterior estimates of the population bias, with a non-significant bias of –0.2%. C. Individual subject results fitted with Bayesian psychometric functions in black (based on 15,000 posterior estimates, 100 representative posterior estimates in red). Blue dots indicating the physical size changes tested by Quest+ (with darkness of the dot indicating the number of times it was tested). Individual biases cluster around zero (from –2.2% to 1.2%). For each participant, alpha (a) is the bias of the logistic function, and beta (b) is the slope.

We fit each of the individual sets of data with a four-parameter logistic Bayesian psychometric function which estimates the slope, the bias, the floor, and the ceiling of the psychometric function (indicated with a black line), using the Palamedes Toolbox 1.10.1 (Prins & Kingdom, 2018) with CmdStan 2.22.0, using the toolbox’s standard priors (bias and slope: normal (0,100), upper and lower lapse rates: beta (1,10)), and based on 15,000 posterior estimates (100 posterior estimates illustrated in red). Fig. 7C shows individual biases range from –2.2% to +1.2%, but cluster around 0, as we would expect if vergence is having no effect on participants’ size judgements. Put simply, participants are perceiving the larger stimuli as larger and the smaller stimuli as smaller, despite the changes in vergence.

To estimate the population level psychometric function illustrated in Fig. 4A, we used the Palamedes Toolbox 1.10.1 (Prins & Kingdom, 2018) with CmdStan 2.22.0 to fit a four-parameter logistic hierarchical Bayesian psychometric function, which fits the data with a multilevel model that takes into account the variability of each subject. We used the toolbox’s standard multilevel priors which are documented by Prins & Kingdom (2019) and, based on 15,000 posterior estimates (100 posterior estimates illustrated in red), found a population level bias of –0.219% (95% CI: – 1.82% to 1.39%) and a population level slope of –0.732 (95% CI: –1.07 to 0.378).

The estimate that particularly interests us is the population bias, so in Fig. 4B we provide a probability density function of the 15,000 posterior estimates of the bias. We find no statistically significant bias, and therefore no statistically significant effect of vergence on perceived size. Indeed, the non-significant bias of –0.2% is in the wrong direction for size constancy.

To go beyond the negative claim that we found no statistically significant effect (null hypothesis not rejected) to the positive claim that there is no effect of vergence on perceived size (null hypothesis accepted), we can make two further arguments.

First, from a Bayesian perspective, we can perform a JZS Bayes factor (Rouder et al., 2009). The estimated Bayes factor that we found was 3.99 (±0.03%), which suggests that the data are four times more likely under the null hypothesis (bias = 0) than under the alternative (bias ≠ 0).

Second, from a frequentist perspective, we can perform an inferiority test that tests whether, if there is a vergence size constancy effect, it is at least as large as the smallest effect size of interest (Lakens et al., 2018). You’ll remember, we defined our smallest effect size of interest as the detection threshold for our most sensitive observer (which is 1.43%). Put simply, any vergence size constancy effect that’s smaller than a 1.43% size change won’t be detected by any of our observers. Since we have a directional hypothesis (vergence micropsia should reduce, rather than increase, the apparent size of the target), we specifically tested whether there is a bias > 1.43%. We therefore performed an inferiority test by taking the 90% confidence interval of the population bias in Fig. 4B in the predicted direction, which is 0.96%. Since this is smaller than 1.43% (our smallest effect size of interest), from a frequentist perspective we can conclude that any vergence size constancy effect is effectively equivalent to zero (Lakens et al., 2018).

## Discussion

According to the literature, “it is well known that vergence is a reliable source of depth information for size constancy” (Sperandio et al., 2013). But we find no evidence that vergence makes any contribution to perceived size. To our knowledge, ours is the first study to report a failure of vergence size constancy at near distances. But ours is also the first study that controls for confounding perceptual cues (changes in the retinal image) whilst also controlling for confounding cognitive cues (keeping subjects naïve about changes in absolute distance).

A number of alternative interpretations of our results have been put to us. In Sections 1-6 of the Discussion we explore these alternative explanations. Whilst we cannot definitively rule out these alternative explanations, we will go on to explain why we do not believe that they give the most plausible interpretation of our results.

### 1. Eye Tracking

We specifically chose not to employ eye tracking in our experiment for four reasons: First, Hooge et al. (2019) find that readily available research eye-trackers “are not accurate enough to be used to determine vergence, distance to the binocular fixation point and fixation disparity”, with errors of up to 2.5°. Second, eye tracking was impractical given our use of parallax barriers, making a clear view of both eyes (for the eye-tracker) and the calibration targets (for the observer) impossible. Third, give the participants were fitted with contact lenses to set their accommodation at 33cm, but the display was at 160cm, ordinary techniques for eye tracking calibration (having participants converge on points on the display 160cm away) could not be employed. Fourth, given our thesis that the vergence size constancy literature reflects cognitive influences rather than truly perceptual effects, we were understandably reluctant to introduce any procedures that would inform participants about the mechanisms underpinning the apparatus, or that eye movements were important for the task being performed.

One suggestion is that the absence of eye tracking leaves open the possibility that participants were not converging during the experiment. Whilst this possibility can’t be excluded, we did employ a subjective criterion for binocular fusion, namely whether the participant reported the target going double. Siegel & Duncan (1960) find that for a 3° target (which ours was, on average) the average maximum disparity participants fail to detect is 2.5° or less. So our subjective criterion of binocular fusion should provide comparable accuracy to eye tracking where the 2D gaze literature (Choe et al., 2016; Drewes et al., 2014; Wildenmann & Schaeffel, 2013; Wyatt, 2010), and the 3D gaze literature (Hooge et al., 2019) report errors of a similar magnitude (up to ≈ 2.5°).

Admittedly, there have been attempts to improve eye tracking accuracy. First, since eye tracking errors are caused by changes in luminance affecting pupil size, one approach is to keep luminance fixed. But it’s not clear that controlling luminance is the right approach for our experiment. If we controlled for luminance in our experiment, as the target got larger it would have to get darker, and as the target got smaller it would have to get brighter. Second, Drewes et al. (2014) found that even if luminance is not controlled, the error can be reduced (down from 2.5° to 0.5°) by calibrating the eye tracker at different luminances. However, this still fails to control for non-luminance effects on pupil size, including changes in pupil size with vergence (the ‘near triad’: Balaban et al., 2018), and cognitive processes (Naber & Nakayama, 2013), although these are liable to be smaller than the errors induced by changes in luminance.

Another concern with our experiment is that participants might have been doing the task with one eye closed, although they were told to keep both eyes open. But both this suggestion, and the suggestion that participants failed to effectively converge, have to be understood in the context of our two other experiments (reported in Linton, 2020) where a similar paradigm was used to demonstrate that vergence was ineffective as an absolute distance cue. For these concerns to be really driving our results in this paper and Linton (2020) we would have to conclude that 35 participants over 3 experiments collectively failed to report pervasive diplopia, or all conducted the experiment with one eye closed, and there is no reason to believe that this is the case.

### 2. Vergence-Accommodation Conflict

Another concern is that vergence and accommodation were decoupled, and this might have affected the vergence response and/or placed the distance estimates from vergence and accommodation in conflict. However, as we have already discussed, some of the most impressive reports of vergence micropsia (that is, vergence affecting the perceived size of objects) occur in the context of pervasive vergence-accommodation conflict (e.g. the 6.5 dioptres of vergence-accommodation conflict in Regan et al., 1986). In this context, +/− 1 dioptres of vergence-accommodation conflict, which is well within the zone of ‘clear single binocular vision’ (Hoffman et al., 2008), should be entirely permissible. Indeed, it is worth considering that 3 out of the 4 contexts I outline in the Introduction in which vergence micropsia has been historically observed (the wallpaper illusion, stereoscopic viewing, telestereoscopic viewing) all rely on inducing much larger vergence / accommodation conflicts than my experiment.

### 3. Vergence vs. Looming

Another suggestion that has been put to us is that our results only pertain to objects looming towards us, either because there is something special about looming, or because the visual system in general is task dependent. It is true that my experiment explicitly introduces a physical size change component into the stimulus. Because the stimulus is viewed in darkness and in isolation, the stimulus appears to move towards the observer when it grows in size, and recede from the observer when it reduces in size. This is well documented for objects changing in size when viewed in darkness and in isolation without a change in vergence (González et al., 2010).

The argument would have to be that changes in the retinal size are so powerful that they eradicate a vergence size constancy signal that is otherwise effective. But then we have to ask what these other contexts might be. The claim of vergence size constancy is that as an object moves towards us in depth in the real world, the visual system cancels out, at least to some extent, its increasing retinal size. But an object moving towards us at near distances in the real world is almost always accompanied by a change in the retinal image size. So in real world viewing, the vergence size constancy mechanism will almost always be accompanied by a looming cue.

After-images are an exception to this rule. But in any case my experiment tests something very close to the after-image literature (Taylor, 1941; Gregory et al., 1959; Carey & Allan, 1996; Mon-Williams et al., 1997; Ramsay et al., 2007; Sperandio et al., 2013). Although Quest+ never tests zero physical size change, as Fig. 7C demonstrates it did test small angular size changes (1-2% over 5 seconds), which are well below threshold for most observers. So, much like an after-image, vergence is changing whilst the retinal image size remains virtually fixed. And yet the points that test these small, non-discriminable, size changes demonstrate that participants are unable to determine whether the target is increasing or reducing in size.

### 4. Gradual Changes in Vergence

Another concern relates to the gradual nature of the vergence changes. Perhaps vergence doesn’t affect perceived size when it is varied gradually (from 50cm to 25cm over 5 seconds), but it is an effective cue to perceived size when varied more rapidly. Again, we cannot rule out this possibility, but would make four observations.

First, we invite readers to try it for themselves. Hold your hand out at 50cm and move it towards your face at 25cm whilst counting out “one thousand, two thousand, three thousand, four thousand, five thousand”. Whilst the motion in depth is gradual, it is very far from being imperceptible, and we would be concerned if a supposedly important perceptual mechanism were unable to process this very apparent motion in depth.

Second, we chose this vergence change because it is exactly the kind of vergence change that had previously been found to produce almost perfect size constancy in Sperandio et al. (2013). Whilst it is open to someone to therefore interpret my results as just a critique of Sperandio et al. (2013), we think this would be a mistake. Since Sperandio et al. (2013) found close to perfect size constancy, if vergence size constancy is ineffective for such gradual changes, then a new (and I would argue cognitive) account must still be posited to explain the close to perfect size constancy in Sperandio et al. (2013). But if this new account can explain the close to perfect size constancy reported in Sperandio et al. (2013), there is no reason why it couldn’t equally explain the close to perfect size constancy reported in other experiments with more rapid eye movements.

Third, you might object that such gradual changes in distance are unusual in the real world. That may be true for an object moving towards us at 5cm/s from 50cm to 25cm. But consider the case where we we’re sitting at a desk looking at a screen. We’re constantly shifting our position in a chair as we look at a screen, moving backwards and forwards towards the screen at a rate of 5cm/s or even less. So what initially looks like an artificial scenario is actually highly relevant to our everyday viewing.

Fourth, notice that this interpretation of our results makes a very specific prediction. Either (a) vergence is not an important cue to size constancy in full cue conditions (a conclusion which I assume those advancing this interpretation would want to resist) or (b) if I gradually move an object in full cue conditions from 50cm to 25cm, observers should misjudge its size because the gradual change in vergence has removed an important size cue.

Note that this final point is the size version of the argument I made in Linton (2020). There I showed that if vergence was manipulated gradually, participants were unable to point to the distance of a target. To maintain that vergence is nonetheless an important absolute distance cue in full cue conditions, one would have to maintain that if I gradually changed the distance of an object in full cue conditions (in the case of Linton, 2020, between 20cm and 50cm), participants should be severely compromised in their ability to point to the distance of the object. If they are not, then it is hard to maintain that vergence is both ineffective as a distance cue when gradually manipulated, and yet an important distance cue in full cue conditions. As I note in Linton (2020), “this account risks replacing the ineffectiveness of vergence under my account with the redundancy of vergence under their account.”

### 5. Limited Changes in Vergence

Another concern is that the eye movements in this experiment are just 25cm in depth, and perhaps vergence size constancy is more effective for larger eye movements. But this concern has to be understood in light of the way in which the vergence angle falls off with distance.

As Fig. 8 demonstrates, assuming 25cm is as close as we look in everyday viewing, then a vergence change from 25cm to 50cm corresponds to around half the full vergence range from 25cm to infinity. If vergence size constancy is not effective in this context, then it cannot be an effective in other contexts (e.g. looking from 50cm to 2m) where the change in distance is much greater, but the change in the vergence angle is much less.

**Figure 8.**
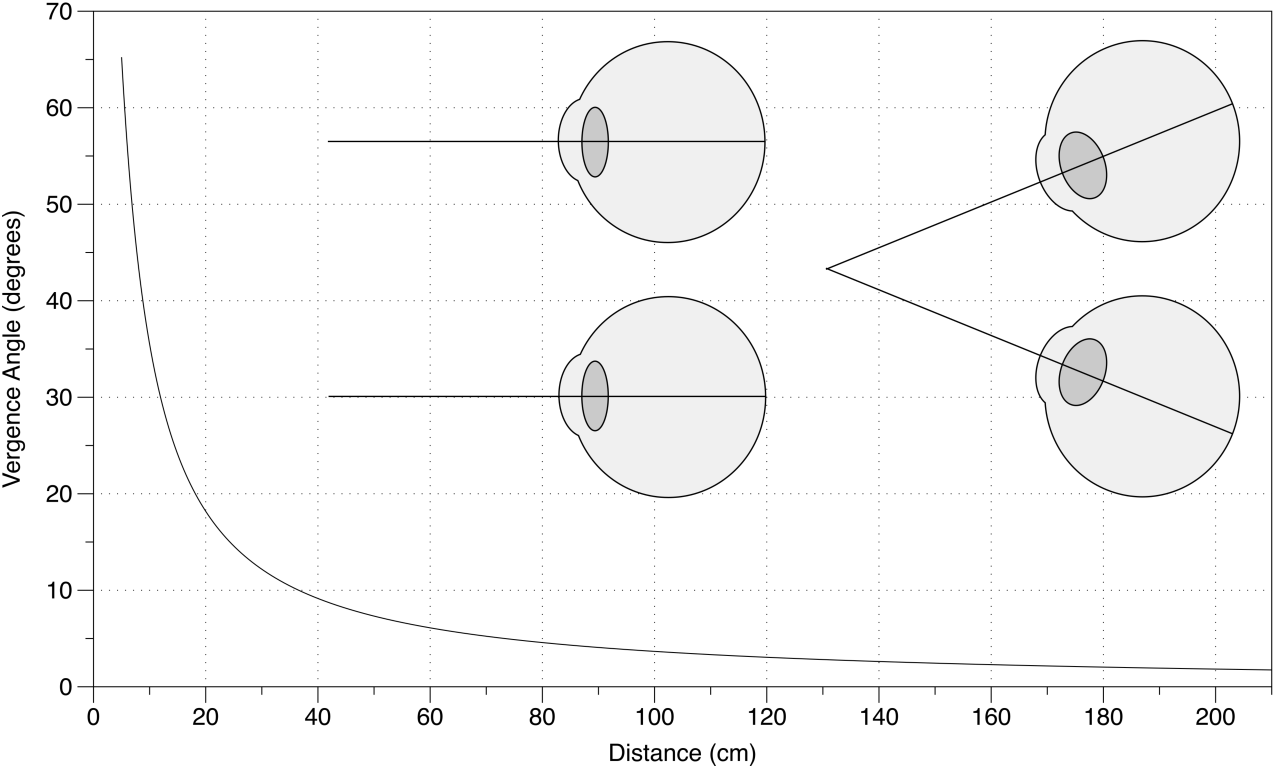
Fall off of vergence angle with distance from Linton (2020). Just as with disparity, the vergence angle falls off as a function of 1/distance^2^.

### 6. Convergent vs Divergent Eye Movements

Since we only tested convergent eye movements (eye movements from far to near), rather than divergent eye movements (eye movements from near to far), conceptually there is the possibility that vergence size constancy still works in one direction (near to far) but not the other (far to near).

However, first, there is no evidence that this is in fact the case. And second, any visual system that increases perceived size when the eyes move from near to far, but doesn’t reduce perceived size when the eyes move from far to near, is going to lead to an ever ratchetting increase in the perceived size of objects as the eyes are moved forwards and backwards in depth. Readers can confirm for themselves that this is not the case.

### 7. Vergence Micropsia

If vergence does not affect perceived size, how do I explain the four contexts where vergence micropsia has been reported in the literature, namely 1. the wallpaper illusion (where we cross our eyes when looking at a wallpaper pattern), 2. stereoscopic viewing (where we vary the vergence whilst looking at two identical images in a stereoscope), 3. telestereoscopic viewing (where we effectively increase the interpupillary distance using mirrors), and 4. the Taylor illusion (where an after-image of the hand appears to shrink when we move it towards us).

There are two explanations of the wallpaper illusion / stereoscopic viewing. First, the same confounding cues I sought to exclude in this experiment (retinal slip and subjective knowledge about our own eye movements) could be used to cognitively infer a reduced size even if this isn’t actually perceived. Second, and more obviously, however, is the fact that the wallpaper illusion and mirror stereoscopes do not control for changes in angular size when we look at a fronto-parallel surface obliquely. This is illustrated by Fig. 5 above (and was controlled for in our experiment using OpenGL). Linton (2018) applies this thought to explain vergence micropsia.

If you cross-fuse the two coins in Fig. 9, the central fused coin appears smaller than when we look at the coins normally. But why does the coin shrink in size if vergence micropsia does not exist? To answer this, we have to understand how the retinal image changes when we cross-fuse.

**Figure 9.**
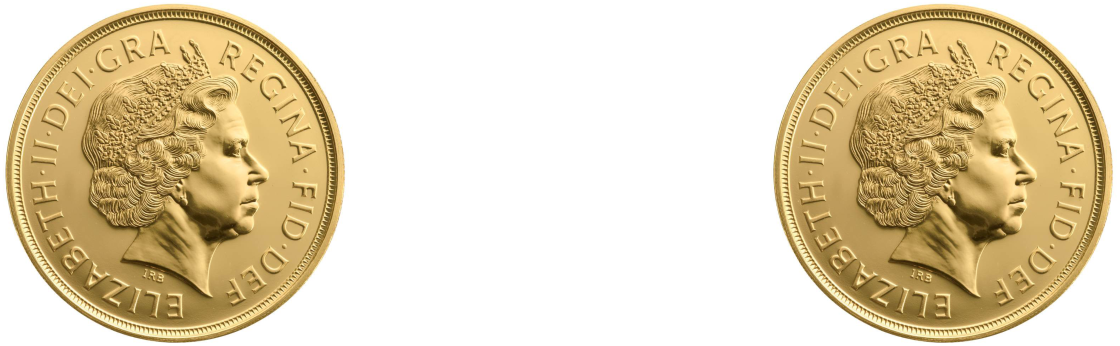
Arrange the coins so they are the interpupillary distance apart (roughly 6cm). Cross fuse the two coins so the right eye looks at the left coin and the left eye looks at the right coin. The central fused coin appears to shrink in size. This is vergence micropsia.

As Fig. 10 illustrates, when we look at the coins normally the right coin projects a larger retinal image to the right eye than the left eye. Equally the left coin projects a larger retinal image to the left eye than the right eye. Now when we cross-fuse, the fused coin is made up of the smaller retinal image of the left coin in the right eye and the smaller retinal image of the right coin in the left eye, explaining why the fused coin is perceived as reduced in scale.

**Figure 10.**
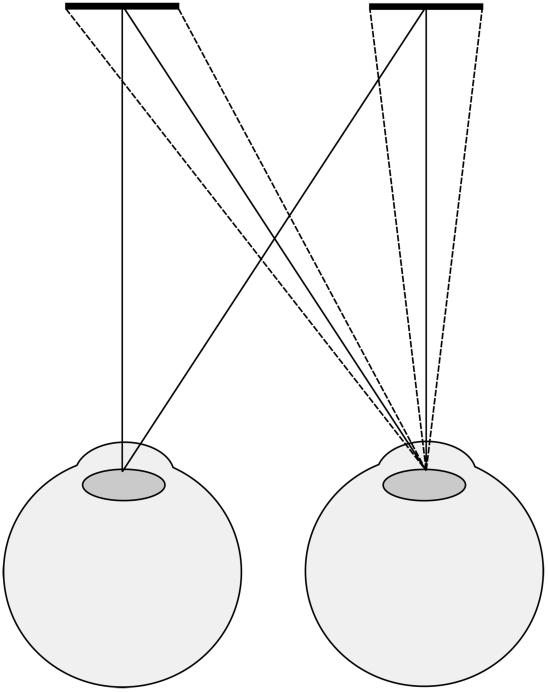
Differences in retinal projections for the left and rights coins in the right eye.

There’s an easy test to check whether this is the correct explanation. This account predicts that the flanking monocular coins either side of the fused coin will be made up of the large retinal image of the left coin in the left eye and the large retinal image of the right coin in the right eye, so the central fused coin should look smaller than the two monocular flankers. At close viewing distances this difference in size between the fused and flanking coins is immediately apparent.

### 8. Cognitive Explanation of Vergence Size Constancy

Let us now turn to the Taylor Illusion as a useful context in which to explore an alternative, purely cognitive, explanation of vergence size constancy. The Taylor illusion (where an after-image of the hand appears to shrink or grow with physical hand movements) is an important paradigm for recent discussions of multisensory integration (Faivre et al., 2017a; Grove et al., 2019). The best current explanation for the Taylor illusion is that it is due (Taylor, 1941; Morrison & Whiteside, 1984; Mon-Williams et al., 1997) or almost entirely due (Sperandio et al., 2013) to the change in vergence as the eyes track the hand moving in darkness. However, in light of our results we suggest that this explanation no longer seems sustainable, since vergence had no effect on the perceived size of the target once subjective knowledge about the fixation distance had been controlled for.

Nor does this imply that the Taylor illusion is primarily due to proprioceptive information from hand motion directly influencing visual perception (Carey & Allan, 1996; Ramsay et al., 2007), since Sperandio et al. (2013) demonstrate that when vergence and hand motion are in conflict, the Taylor illusion follows vergence, and the effect is only marginally reduced in size.

Instead, what both accounts are missing is the participant’s subjective knowledge about their own changing hand and gaze positions. We would suggest that explains why Sperandio et al. (2013) found that vergence affects perceived size when their participants knew about their changing gaze position (from their hand or from the motion in depth of an LED), but why we didn’t when our participants were ignorant of this fact. There are two ways in which conscious knowledge about our changing hand or gaze position could influence size constancy.

First, our subjective knowledge could influence our visual experience (so-called ‘cognitive penetration’ of perception by cognition). But we are skeptical of invoking ‘cognitive penetration’ to explain an effect that could also be explained as a purely cognitive bias (for further skeptical discussions of ‘cognitive penetration’ see Fodor, 1983; Pylyshyn, 1999; Firestone & Scholl, 2016).

Second, under our alternative cognitive bias account, the integration of the retinal image and our changing gaze position could be purely cognitive, rather than perceptual. Our visual experience of the after-image’s angular size remains constant, but because we know that our hand is moving towards our face, our hand movements cognitively bias our interpretation of our constant visual experience and we interpret the constant angular size of the after-image as a reduction in physical size.

Put another way, the after-image literature can’t place itself outside the looming literature. The absence of a change in retinal image size is still a looming cue. The fact that after-images don’t change in retinal size, whilst the participant knows that they change in distance (for instance, due to retinal slip from diplopia, or from motion of their own hand in the Taylor illusion) is a cue that the object is shrinking in size, because there is an absence of the attendant increase in retinal size that one would expect. This is an entirely consistent looming-based explanation of the vergence size constancy after-image literature that does not invoke the notion of vergence size constancy.

One reviewer has asked why subjective knowledge should determine perceived size in this context but not e.g. in the Ames Room where we know that the two people we are seeing are the same size. This is an open question, but note two disanalogies between these two scenarios. First, the Ames Room is about perceived physical size (judging the wrong physical size given the different angular sizes of the people), whilst the Taylor illusion is primarily about perceived angular size (the impression of the hand ‘shrinking’ or ‘growing’). Second, the Taylor illusion occurs over time, and without the opportunity for direct comparison between the target at t_1_ and t_2_, implying an important role for working memory, and therefore additional scope for cognitive influences.

### 9. Multisensory Integration

This purely cognitive interpretation of the Taylor illusion has wide reaching implications for multisensory integration, specifically the integration of vision and hand movements. The Taylor illusion is taken as evidence of multisensory integration at the level of perception. Specifically, that vision “relies on multimodal signals” (Sperandio et al., 2013; Chen et al., 2018) and that “visual consciousness is shaped by the body” (Faivre et al., 2015; Faivre et al., 2017a; Faivre et al., 2017b). But if the integration of proprioception and the retinal image could be purely cognitive in the context of vergence (the major driver of the Taylor illusion in Sperandio et al., 2013), there’s no reason why the integration of proprioception and the retinal image in the context of integrating vision and hand movements (the minor driver of the Taylor illusion in Sperandio et al., 2013) couldn’t equally be accounted for in purely cognitive terms.

This cognitive approach also suggests a non-perceptual explanation for variants of the Taylor illusion that appear to demonstrate the integration of vision with the rubber-hand illusion (Faivre et al., 2017a) and tool use (Grove et al., 2019).

And cognitive interpretations of the integration of vision and proprioception are also advanced in the contexts of vision and touch in slant estimation (Hillis et al., 2002; Gepshtein et al., 2005) by Linton (2017), pp.37-38 and pp.65-66, and vision and vestibular cues in self-motion (Fischer & Kornmüller, 1930; Ash et al., 2011) by Linton (2018).

### 10. Vergence Modulation of V1

Ever since Trotter et al. (1992) found that the large majority of neurons in the monkey primary visual context (V1) were modulated by vergence, it has been suggested that processing of the vergence signal in V1 plays an important role in size constancy. Further evidence for the vergence modulation of V1 is found by Trotter et al. (1993); Trotter et al. (1996); Dobbins et al. (1998); Trotter & Celebrini (1999); Cumming & Parker (1999); Trotter et al. (2004); Cottereau et al. (2014). Cumming & Parker (1999) found a smaller proportion of neurons in V1 were affected, and to a less dramatic extent, than in Trotter et al. (1992; 1996). One reason could be the differences in vergence distances tested, but another possibility raised by Cumming & Parker (1999) is that since vergence was not tracked in Trotter et al. (1992; 1996), there’s no guarantee that poor fixation didn’t cause retinal disparities. This concern might also apply to other studies.

Related discussions include the role of vergence in the parietal cortex (Gnadt & Mays, 1991; 1995; Quinlan & Culham, 2007; Culham et al., 2008), and potentially also LGN (Richards, 1968; although this was speculative, and never followed up), as well as early neural network models (Lehky et al., 1990; Pouget & Sejnowski, 1994) that Trotter et al. (1992) complements.

More recently Chen et al. (2019) have looked at the time course of size constancy, and found that vergence and the retinal image are not integrated during (a) initial processing in V1 (∼50ms), but instead during (b) recurrent processing within V1, and/or (c) re-entrant projections from higher-order visual areas (e.g. Gnadt & Mays, 1991; 1995), both of which are consistent with the ∼150ms timeframe. This is consistent with Trotter et al. (1992)’s suggestion that whilst vergence responsive neurons encode vergence distance, further computations are required to scale the retinal image, so vergence responsive neurons “constitute an intermediate step in the computation of true depth, as suggested by neural network models [Lehky et al., 1990].”

However this whole line of research, from Trotter et al. (1992) to the present, is prefaced on the fact that “psychophysical data suggest an important role for vergence” (Trotter et al., 1992). And this is what our results in this experiment, and in Linton (2020), question. Taken together, our results in this paper and in Linton (2020) suggest that there is no link between vergence and distance perception (Linton, 2020) or size perception (this paper), and therefore no link between the vergence modulation in V1 (or anywhere else) and distance or size perception.

How, then, are we to account for the vergence modulation of V1 that has been reported by Trotter et al. (1992; 1996). There are three possibilities:

First, one objection to our account is that we have not excluded every possible task in which vergence might be implicated in size and distance perception. One possibility that is often suggested is that distance is extracted from vergence in order to scale binocular disparities for our perception of 3D shape (Wallach & Zuckerman, 1963), and that errors in this mechanism are responsible for distortions of visual shape (Johnston, 1991). This will be the focus of future experimental work, although we would be surprised if distance from vergence was used to scale binocular disparities whilst failing to scale size and distance themselves.

Second, as we have argued throughout this paper, we should be careful attributing effects to vergence that could equally be explained by changes in the retinal image. Cumming & Parker (1999) raise the prospect of vergence insufficiency leading to changes in retinal disparity. Another possibility is that the results in Trotter et al. (1992; 1996) simply reflect changes in vertical disparity with distance, since Trotter et al. (1992; 1996) change the distance of the display, not vergence per se. So the vergence modulations in V1 might simply be artifacts.

Third, according to our alternative account (Linton, 2018a), visual scale is entirely dependent upon top-down cognitive processing. But as Linton (2021) argues, this approach is entirely consistent with evidence of eye movement processing in V1, even if this processing simply reflects subjective knowledge about our changing gaze position. There is increasing evidence that V1 is implicated in purely cognitive (non-visual) processing, e.g. the location of rewards amongst visually identical targets (Saleem et al., 2018). So the vergence modulations of V1, if they do exist, could equally reflect the participant’s own purely cognitive knowledge about their changing gaze position, even though this has no effect on their visual experience.

### 11. Telestereoscopic Viewing

Of the four instances of vergence size constancy we discussed in the Introduction, alternative explanations of (a) the wallpaper illusion and (b) stereoscopic viewing have been provided in Section 7, whilst an alternative explanation of (c) the Taylor illusion is provided in Section 8. That only leaves (d) telestereoscopic viewing to be explained. This is when the interpupillary distance of the eyes is increased using mirrors, leading to the impression that the world has been ‘miniaturised’. Helmholtz (1857; 1858; 1866, p.310) explained this miniaturisation in terms of the increase in vergence required to fixate on an object (Fig. 11).

**Figure 11.**
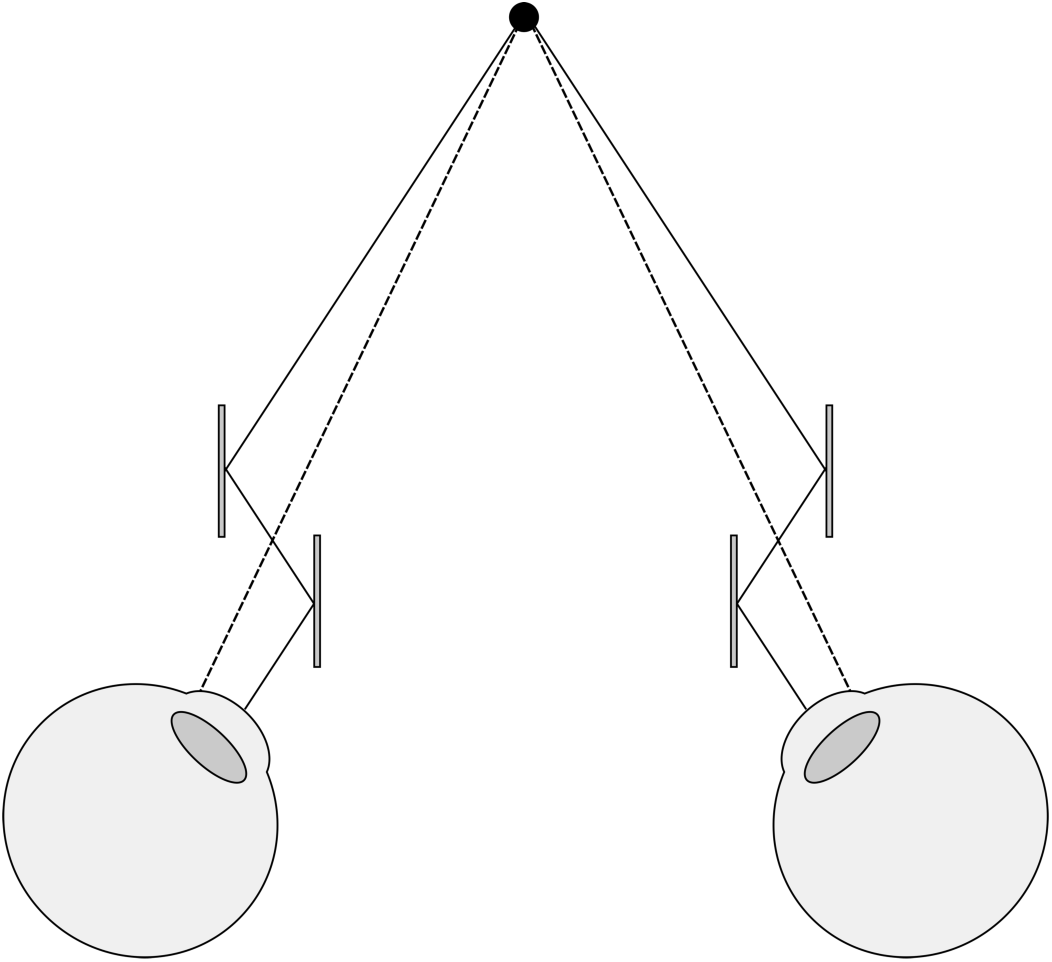
Telestereoscopic viewing. Mirrors alter the path of light to the eyes. In order to view the same point in space, the eyes now have to rotate more (as indicated by the solid line) than they ordinarily would (as indicated by the dotted line).

However, our results in this paper challenge this account.

An alternative explanation for this effect, and potentially the micropsia effects in the wallpaper illusion and stereoscopic viewing (although not the Taylor illusion), are changes in vertical disparities. Although Rogers (2009; 2011) appeals to vertical disparities as well as vergence to explain telestereoscopic viewing, we are skeptical of this explanation.

The initial promise of vertical disparities was that you could take any three points in the scene (and potentially just two), and determine the geometry and viewing distance of the scene (Longuet-Higgins, 1982) (see also Mayhew, 1982; Mayhew & Longuet-Higgins, 1982). However, there’s no evidence the visual system can achieve this with such a limited number of points, and the emphasis soon shifted (in Gillam & Lawergren, 1983) to the gradient of vertical disparities across a fronto-parallel surface. Rogers (2017) explains the contemporary vertical disparity literature in the following terms: “For any continuous surface like the chessboard, there is a horizontal gradient of the ratio of the vertical sizes in the two eyes and this varies with viewing distance.” But there are four concerns with account:

First, telestereoscopic viewing should not affect our perception of size and distance under the classic Longuet-Higgins (1982) account, since in the process of estimating egocentric distance this account also estimates the interpupillary distance and vergence angle. Put simply, whether our eyes are far apart and rotated a lot, or close together and rotated a little, so long as they are fixated on the same point in physical space, the approach in Longuet-Higgins (1982) should give exactly the same egocentric distance, and therefore exactly the same perceived scale.

Second, vertical disparities have proved an ineffective absolute distance cue for surfaces that take up as much as 25° x 30° of the visual field (Cumming et al., 1991; Sobel & Collett, 1991). Rogers & Bradshaw (1995) argue that this is because vertical disparities are maximised in the periphery. But even the results for a 60° surface are questionable. Vienne et al. (2016) investigated how vertical disparities could improve the distance perception of a cylinder in front of a 60° background. However, first, subjects performed no better, and potentially performed worse, with the 60° background present than without it. Second, whilst increasing the vertical disparities of the background did change the apparent distance of the cylinder, this was largely because participants changed their previous judgements about the cylinder’s distance when the background’s vertical disparities were undistorted. Had participants made consistent judgements about the same stimulus across conditions the effect would largely disappear. If there is no benefit of having a 60° background, and no consistency in distance judgements across conditions when it is present, it becomes hard to maintain that vertical disparities provide an effective absolute distance cue.

Similarly, it is hard to claim that Rogers & Bradshaw (1995) show more than an ordinal effect of vertical disparities on distance judgements even for a 70° surface, since “different observers differed in the way they used numbers to indicate absolute distances”, and so distances underwent a normalisation and scaling process.

Third, even taking Vienne et al. (2016) (60°) and Rogers & Bradshaw (1995) (70°) results at face value, what do they imply? Taken on their face value these results limit the application of vertical disparities to (1) regularly textured flat surfaces, that are (2) no further away than 1-2m (see the sharp fall-off of vertical disparities with distance for a 60° x 60° surface in Vienne et al., 2016, Fig. 1c), and (3) take up at least 30° (and arguably 60°-70°) of the visual field. Since such surfaces are almost never encountered in natural viewing conditions, this cannot be how we experience visual scale. Theoretically the results in Rogers & Bradshaw (1995) and Vienne et al. (2016) could support a broader principle, but as Rogers & Bradshaw (1995) note: “Whether the visual system extracts the vertical disparities of individual points or the gradient of VSRs over a surface is an empirical question for which there is no clear answer at present.”

Fourth, even if a fronto-parallel surface is not required, the points still need to take up 60°-70° of the visual field. This can’t explain how telestereoscopic viewing can change the perceived scale of a photograph which takes up much less than 60°-70° when viewed. See for example Fig. 12 and Fig. 13. Looking at them through red-blue glasses with one eye open, they look like normal photos. But as soon as you view them through red-blue glasses with both eyes open, you get the immediate impression that we are “not looking at the natural landscape itself, but a very exquisite and exact model of it, reduced in scale” (Helmholtz, 1866, p.312). Whatever explains the miniaturisation effect of telestereoscopic photographs likely explains the miniaturisation effect of telestereoscopic viewing in the real world, and yet large angular sizes are not required for telestereoscopic photographs to be effective.

**Figure 12.**
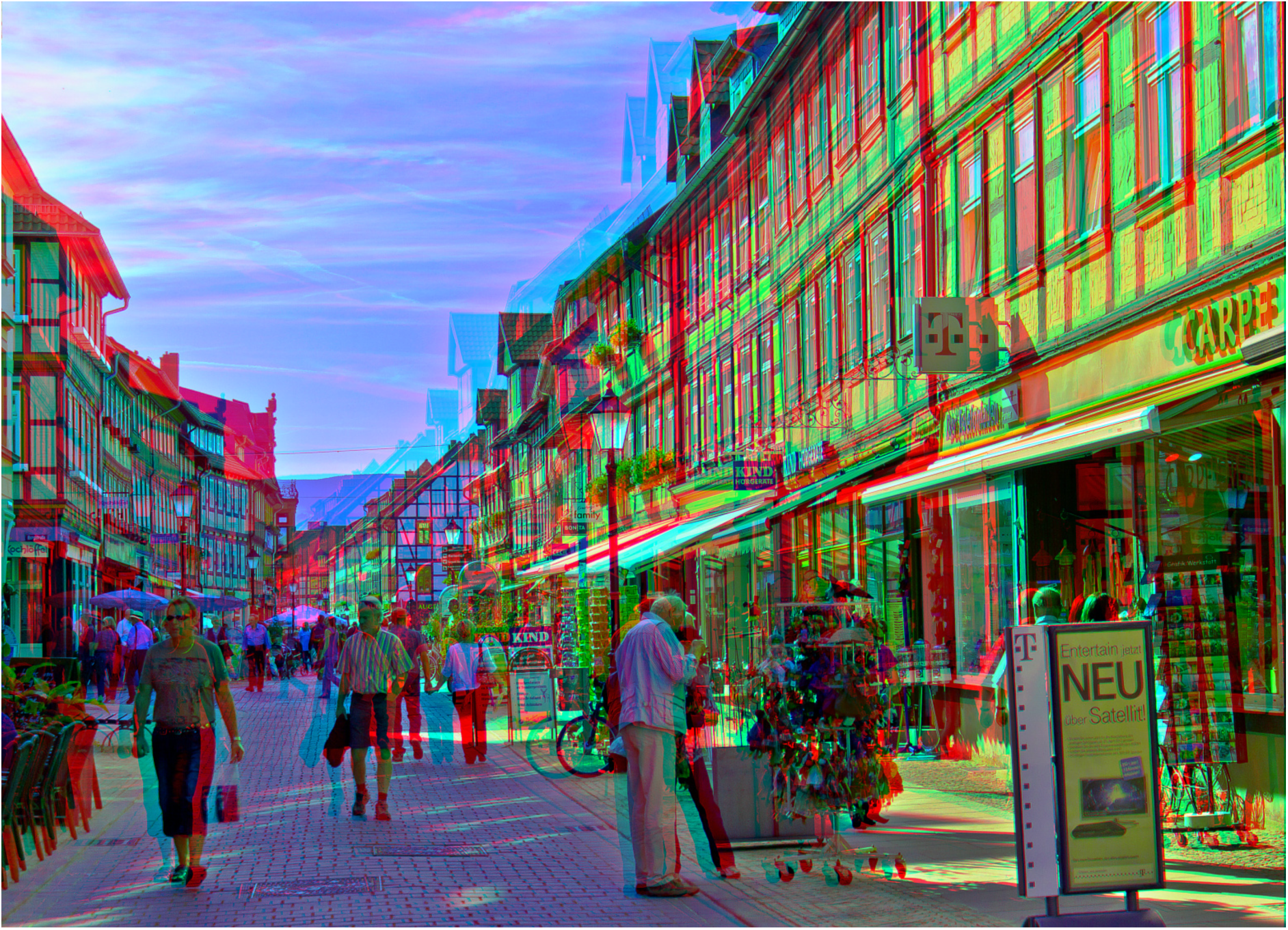
‘Wernigerode Boulevard’ (2011) by Sasha Becher. © Sasha Becher. Used with permission from: https://www.flickr.com/photos/stereotron/6597314627/in/album-72157612377392630/ For more of Sasha Becher’s telestereoscopic images please see: https://www.flickr.com/photos/stereotron/

**Figure 13.**
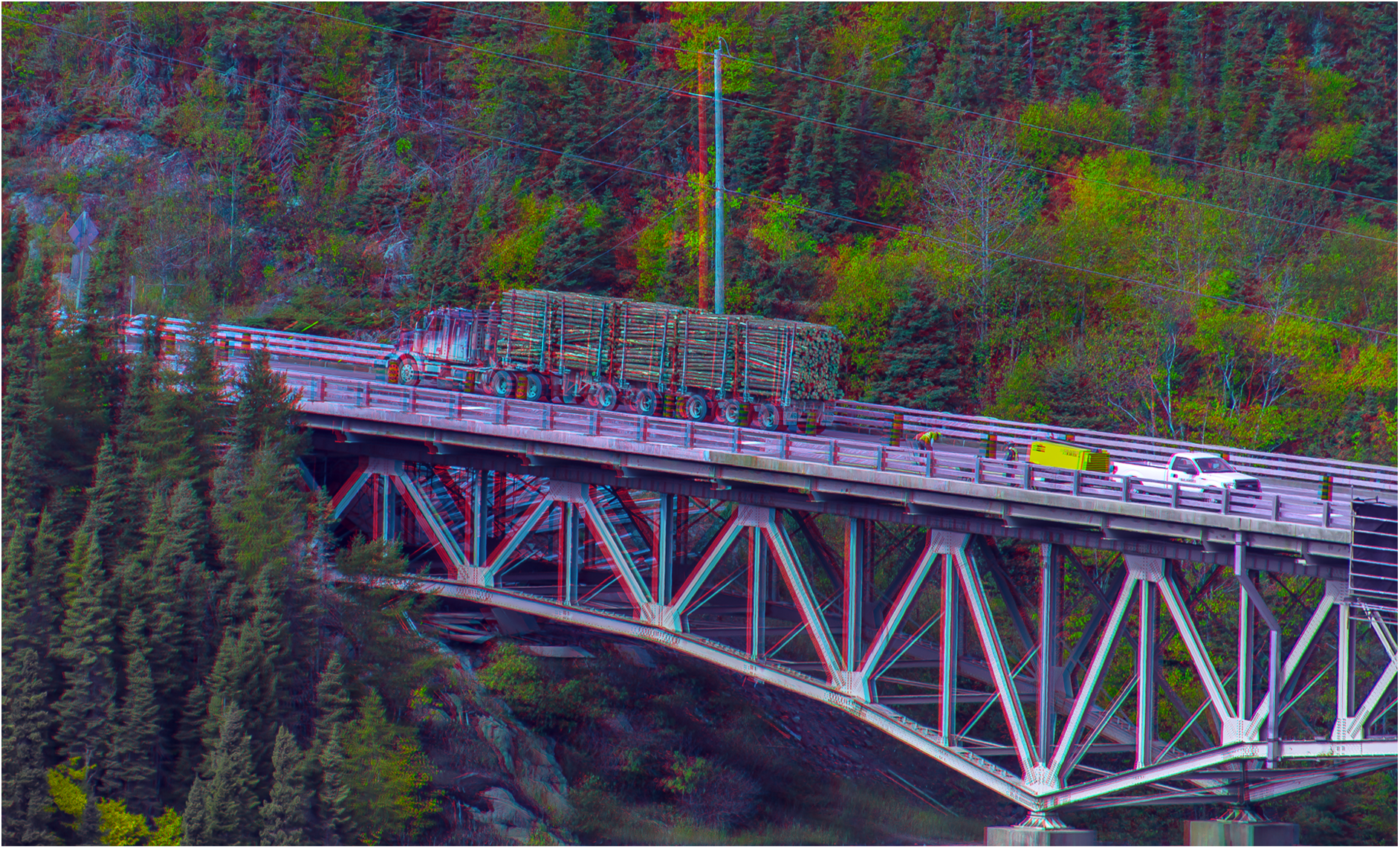
‘Trans-Canada Highway @ Neyes National Park’ (2014) by Sasha Becher. © Sasha Becher. Used with permission from: https://www.flickr.com/photos/stereotron/28817920100/in/album-72157612377392630/ For more of Sasha Becher’s telestereoscopic images please see: https://www.flickr.com/photos/stereotron/

Instead of vergence or vertical disparities, in Linton (2018a) I advance a third distinct explanation for the change in scale experienced in telestereoscopic viewing based on horizontal disparities. The argument begins from the premise that this is the only remaining parameter that is manipulated in telestereoscopic viewing apart from vergence and vertical disparities. Horizontal disparities are typically thought of merely as an affine depth cue (not even a relative depth cue) until they are scaled by an absolute distance cue, typically thought to be vergence (Wallach & Zuckerman, 1963; Johnston, 1991). The reason for this is that binocular disparities fall off with 1/distance^2^, and so in order to extract relative depth (which is a relationship in terms of 1/distance not 1/distance^2^) we need to know the viewing distance. So, on this account, distance from vergence provides us with perceived stereo depth (Johnston, 1991): ‘Distance → Stereo Depth’. My argument in (Linton, 2018a) seeks to invert this relationship (known as ‘depth constancy’), to instead go from ‘Stereo Depth → Distance’, in three stages:

First, on this account, there is no scaling of binocular disparity using vergence, and therefore no ‘depth constancy’. Whilst Johnston (1991) found some evidence of ‘depth constancy’, this was not tested in the controlled blackout conditions of Linton (2020) or this paper. Under this alternative account, perceived depth simply reflects binocular disparity that falls off with 1/distance^2^. This will be the subject of future empirical investigation.

Second, under this account, we are unconsciously very sensitive to the way in which perceived depth falls off with distance. What I describe as “a cognitive association between (a) vivid stereo depth and (b) closer distances (reflecting our experience of an environment where disparity falls-off with distance^2^)” in Linton (2018a). Put simply, we only ever experience the vivid stereo depth we experience in Figures 12 and 13 when we view a scene up close, and so we interpret it as miniature (Linton, 2020b).

Whilst vivid stereo depth equates to near distances, flat stereo depth could equate to ‘far away’ or ‘up close and flat’, so assumptions about scene geometry will often be necessary. Our interpretation of the relationship between stereo depth and distance is informed by natural scene statistics (Linton, 2020b), enabling us to make appropriate assumptions.

The relationship between stereo depth and distance would be more informative if we could compare the perceived geometry of the scene (depth from disparity) with the actual geometry of the scene. So, in Figures 12 and 13, we might additionally infer that the accentuated stereo-depth in Figures 12 and 13 is only consistent with the geometry of the scene if the scene is being viewed up close; i.e., if the scene is miniature (Linton, 2020b). If we do, it may be that we recognise the scene as of a certain familiar kind. Or it may be that we rely on pictorial cues, which are comparatively distance invariant (albeit subject to perspective distortions / foreshortening: Tehrani et al., 2016), and which are unaffected by telestereoscopic viewing (which increases binocular disparity without reducing viewing distance).

I would expect the same miniaturisation effect to be present if the scenes in Figures 12 and 13 were depicted using a random-dot stereogram. This doesn’t necessarily rule out familiarity or pictorial cues. In Linton (2017) we argue that the cues that inform us about scene geometry in a pictorial sense are processed after stereo depth has been perceptually resolved. For instance, you can embed a 2D line drawing of a cube within a random dot stereogram. The line drawing only emerges after the stereo depth has been perceptually resolved. But you can still recognise it as a 2D line drawing of a cube. So you’re interpreting the shape of the cube using pictorial cues, such as perspective, much like any 2D line drawing. But it would be a mistake to suggest that this necessarily implies that what you are relying on are monocular cues because they don’t exist. Similarly, you can imagine viewing a face in a random dot stereogram. The face might be accentuated or reduced in depth, but you’d still be able to understand it as a face, and therefore interpret it as being accentuated or reduced in depth even though there are no monocular cues.

Third, we claim that the relationship between accentuated stereo depth (i.e. accentuated 3D shape) and visual scale is merely a post-perceptual cognitive association. What does this mean? So far as our visual experience is concerned, all that is being manipulated with telestereoscopic viewing is our perception of 3D shape. An illustration of this is provided with Fig. 12 and Fig. 13. Close one eye and we don’t have the apparent miniaturisation in scale that we do with binocular vision. And yet what changes between monocular and binocular viewing? First, the angular size of the objects in the scene does not appear to change in any dramatic sense, so we cannot attribute to the dramatic change in perceived scale to a dramatic change in perceived angular size. Second, the distance of an object in the image does not appear to change dramatically when we fixate on it and close one eye. So there is no sense that our perception of (a) size or (b) distance changes dramatically as we fixate on an object in the scene and open and close one eye. Instead, the defining change in our visual experience as is the change in the perceived 3D shape of the scene.

Note that it might be tempting to write off telestereoscopic viewing as a contrived illusion. But that would be a mistake. What telestereoscopic viewing demonstrates is that binocular cues dominate all other cues to absolute size and distance. In telestereoscopic viewing all other cues to absolute distance in the real world are maintained, and yet our impression of visual scale varies dramatically with the manipulation of the inter-pupillary distance using mirrors. Instead of interpreting our inter-pupillary distance as changing, we attribute this change to the scene itself, and see it as miniaturised. Whatever explains why we see the world as the wrong size when binocular cues are wrong (in telestereoscopic viewing) also explains why we see the world as the right size when binocular cues are correct (in ordinary binocular viewing).

If this is the correct interpretation of telestereoscopic viewing, then our results suggests that visual scale is much more reliant on cognitive influences than previously thought. First, because our results challenge the suggestion that size and distance are triangulated using vergence. Second, because our alternative explanation of binocular scale perception in telestereoscopic viewing is entirely cognitive in nature. Our results are consistent with our argument in Linton (2017; 2018) that visual scale is based solely on higher level cognitive processes, where we extend Gogel (1969)’s and Predebon (1992)’s observations about familiar size to argue that visual scale itself is a purely cognitive process.

### 12. Development of Size Constancy

One objection to our account is that infants 1.5-2 months of age can apparently respond to the physical, and not just the retinal, size of objects (Bower, 1965; Granrud, 1987; Slater et al., 1990). According to this objection, this speaks against a cognitive explanation. However, we have good reason to be cautious of the claims in this literature. First, Bower (1965) found no benefit of binocular vision on infant size constancy, grounding his account in motion parallax instead. This should immediately alert us that this literature is missing something important. Second, even on motion parallax, Bower admitted that infants could simply be responding to changes in the stimulus with head movements, rather than perceived size. Third, a counterpoint to this literature is that those with their sight restored after early blindness often experience gross errors in their distance judgements (Gregory & Wallace, 1963; Šikl et al., 2013). Fourth, whilst it is merely anecdotal, Helmholtz (1866)’s recollection of his own gross failures of size constancy as a child (Vol.3, p.283) inspired his whole account of inferential size and distance perception.

## Conclusion

Vergence is thought to provide an essential signal for size constancy. We tested vergence size constancy for the first time without confounding cues, and found no evidence that eye movements make any contribution to perceived size. We explore a number of alternative explanations for these results. Whilst we cannot definitively exclude these alternatives, we conclude that the most plausible interpretation of our results is that vergence does not contribute to size perception, complimenting previous results (Linton, 2020) that also suggest that vergence does not contribute to distance perception. If this is the correct interpretation, this work has three important implications. First, it suggests that our impression of visual scale is much more reliant on cognitive processing than previously thought. Second, it leads us to question whether the vergence modulation of neurons in V1 reported in the literature really contributes to our impression of visual scale. Third, it leads us to question whether the multisensory integration reported in the context of the retinal image with proprioceptive cues from the hand is better understood in terms of observers having subjective knowledge about their hand and gaze position.

## Acknowledgements

This research was conducted under the supervision of Christopher Tyler and Simon Grant. We thank Salma Ahmad, Mark Mayhew, and Deanna Taylor, for fitting the contact lenses, and Jugjeet Bansal, Priya Mehta, and the staff of the CitySight clinic for their assistance. We thank Joshua Solomon, Matteo Lisi, Byki Huntjens, Michael Morgan, Chris Hull, and Pete Jones for their advice. We thank audiences at the Applied Vision Association (2019), the British Machine Vision Association’s ‘3D worlds from 2D images in humans and machines’ meeting (2020): https://youtu.be/6P3EYCEn52A and the (Virtual) Vision Sciences Society (2020): https://osf.io/tb3un/ / http://youtu.be/VhpYjPj5Q80 where this work was presented, as well as audiences at the Association for the Scientific Study of Consciousness (2019) where the argument about telestereoscopic viewing was presented: https://osf.io/9ry3s/. We thank Scott Murray for allowing me to reproduce Fig. 1, and Sascha Becher for allowing me to reproduce Fig. 12 and Fig. 13. We also thank the three referees and the two editors for their very valuable comments on the paper.

## Open Practice Statement

Code for running the experiment, and all the data and analysis scripts, are accessible in an open access repository: https://osf.io/5nwaz/

